# Modeling Substrate Coordination to Zn-Bound Angiotensin Converting Enzyme 2

**DOI:** 10.1101/2021.03.27.437352

**Authors:** Peter R. Fatouros, Urmi Roy, Shantanu Sur

**Affiliations:** Department of Chemical and Biomolecular Engineering, Clarkson University, 8 Clarkson Avenue, Potsdam NY 13699, United States; Department of Chemistry and Biomolecular Science, Clarkson University, 8 Clarkson Avenue, Potsdam NY 13699, United States; Department of Biology, Clarkson University, 8 Clarkson Avenue, Potsdam NY 13699, United States

**Keywords:** angiotensin converting enzyme 2, angiotensin II, binding free energy, metalloenzyme, molecular dynamics simulation, zinc parameterization

## Abstract

The spike protein in the envelope of severe acute respiratory syndrome coronavirus 2 (SARS-CoV-2) interacts with the receptor Angiotensin Converting Enzyme 2 (ACE2) on the host cell to facilitate the viral uptake. Angiotensin II (Ang II) peptide, which has a naturally high affinity for ACE2, may be useful in inhibiting this interaction. In this study, we computationally designed several Ang II mutants to find a strong binding sequence to ACE2 receptor and examined the role of ligand substitution in the docking of native as well as mutant Ang II to the ACE2 receptor. The peptide in the ACE2-peptide complex was coordinated to zinc in the ACE2 cleft. Exploratory molecular dynamics (MD) simulations were used to measure the time-based stability of the native and mutant peptides and their receptor complexes. The MD-generated root-mean-square deviation (RMSD) values are mostly similar between the native and seven mutant peptides considered in this work, although the values for free peptides demonstrated higher variation, and often were higher in amplitude than peptides associated with the ACE2 complex. An observed lack of a strong secondary structure in the short peptides is attributed to the latter’s greater flexibility and movement. The strongest binding energies within the ACE2-peptide complexes were observed in the native Ang II and only one of its mutant variants, suggesting ACE2 cleft is designed to provide optimal binding to the native sequence. An examination of the S1 binding site on ACE2 suggests that complex formation alone with these peptides may not be sufficient to allosterically inhibit the binding of SARS-CoV-2 spike proteins. However, it opens up the potential for utilizing AngII-ACE2 binding in the future design of molecular and supramolecular structures to prevent spike protein interaction with the receptor through creation of steric hindrance.

---

COVID-19 pandemic caused by SARS-CoV-2 has emerged as a major health issue with global impacts higher than any other infectious disease in recent history^1^. One major challenge in containing the disease spread stems from a high level of respiratory transmission. Decreasing the ability of the virus to infect the respiratory epithelial cells could offer an effective measure to control disease spread and severity. To infect hosts’ cells, spike (S) protein found in SARS-CoV-2 will first be primed by the transmembrane serine protease 2 (TMPRSS2), acting on the S2 domain.^2,3^ The S1 domain of the primed S protein, displayed on the virus surface, will then interact with angiotensin converting enzyme 2 (ACE2) located in the lipid rafts of hosts’ cell membranes, inducing endocytosis and eventual infection.^4,5^ To prevent the viral infection, several research groups have aimed to block this interaction by inhibiting the S protein.^6–8^ This approach requires the design of novel inhibitors with high affinity for the S protein. Alternatively, inhibitors can be developed for ACE2 utilizing its native substrate, angiotensin II (Ang II) peptide as a starting structure. Physiologically, ACE2 cleaves a single amino acid from the C-terminus of the octapeptide Ang II and helps to regulate blood pressure. ACE2 can be classified as a gluzincin from the amino acid sequence found coordinating to zinc at the Ang II binding site.^9,10^ From the known properties of this class of metalloprotease, the catalysis will occur through nucleophilic addition of a water molecule coordinated to zinc found within the ACE2 structure.^10^ Although the affinity of Ang II for ACE2 is high, it does not share an active binding site with S protein, and thus might not be effective alone in blocking the binding of S protein.^9,11^ However, Ang II peptide can be incorporated in a peptide-nanoparticle conjugate^12^, which could physically block the S protein interaction with ACE2 following binding of Ang II at its receptor site. The success of such approach will highly depend on the affinity of Ang II for ACE2 and the affinity can potentially be improved through alterations in the Ang II primary structure as shown in the study by Clayton et al.^13^ Although this work has experimentally screened a number of mutant Ang II sequences, several other sequence possibilities are yet to be examined. This study aims to use computational methods to determine and compare the affinities of reported and additional unexamined Ang II mutants with the goal of finding a sequence that binds strongly to ACE2.

Computational approaches are commonly used in drug design as they allow for reductions in cost and time when scanning for new targeting molecules. Several groups have used computational strategies to search for new inhibitor molecules for S protein and ACE2.^14,15^ In these approaches, potential inhibitors are docked to the receptor and undergo molecular dynamics (MD) simulations followed by calculations of binding free energy. In these studies, the ability to accurately estimate the interactions that could be further experimentally validated, depend critically on the assignment of proper parameters to all atoms and interactions found within the system for the MD simulations and binding free energy calculations.

Although it is estimated that 10% of the human proteome contains zinc, it remains challenging to completely parameterize it for MD simulations.^16^ Interactions between zinc and amino acids (cysteine, histidine, tyrosine, and carboxylic acids) can be unusually strong and can involve complex charge transfer and polarization, which are not accounted for in classical Amber or CHARMM forcefield parameters.^17–21^ While several methods have been developed to parameterize these interactions, these strategies generally fall into two main categories, namely bonded and non-bonded models. The bonded model generates explicit bonds between zinc and coordinating atoms, fixing the coordination geometry and preventing ligand exchange.^22–25^ This method also involves the calculation and assignment of partial charge on the zinc ion, which is more realistic than assignment of the formal +2 integer charge.^25–27^ In contrast, the non-bonded

model assigns a +2 integer charge on zinc and allows for ligand exchange during MD simulations.^28^ However, it is accepted that this model fails to describe tetra-and penta-coordinated structures and tends to prefer octahedral coordination geometries.^22,29–31^

In this work, we design, dock, and simulate several Ang II mutant sequences to find a strong binding sequence for ACE2. Native peptide and each mutant was docked to zinc at each of the possible coordination sites found within the peptide, replacing the existing water molecule at the coordination site. We also examine the potential impact of peptide binding to ACE2 on the structure of the S protein binding region.

## RESULTS

To study the influence of peptide sequence on ACE2 binding, we considered seven mutant Ang II peptides and compared those against the native Ang II sequence (Figure1A). The mutant peptides were obtained by making single or double mutations at amino acid positions 4-6 of the native sequence. The structures of these peptides are included in the supporting information (SI) Figure S1. In previous experimental work, two of these sequences (DRVYI**Y**PF and DRVY**VY**PF) were demonstrated to increase ACE2 inhibition and reduce peptide cleavage^13^. The other five mutants were designed to test the replacement of tyrosine with the stronger coordinating residues, glutamic acid and cysteine. The stability of the mutant peptides was analyzed by computing the RMSD over a 48 ns production run (Figure 1B, S2). We observed that the average RMSD values and the magnitudes of fluctuations of these mutant peptides (range of average RMSD and standard deviations: 3.96-5.51 ± 0.67-1.10 Å) were comparable to the average RMSD value of native Ang II (4.76 ± 0.90 Å). The relatively high fluctuations in RMSD values can be attributed to the lack of organized secondary structure within these small peptides.

**Figure 1.**
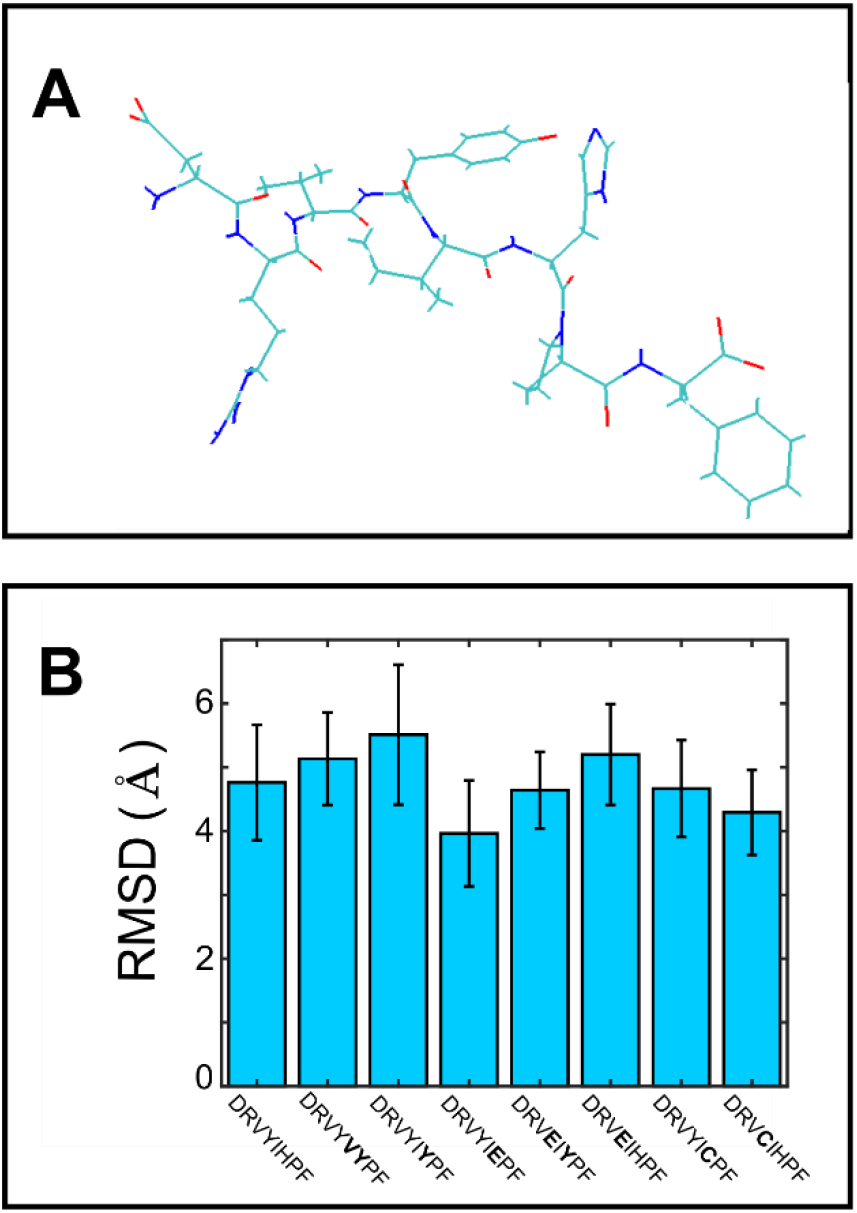
(A) The minimized and equilibrated structure of Ang II. (B) Averaged RMSD of Ang II and seven Ang II mutants during a 48 ns production run. The bolded residues found in the peptide sequences indicate the mutations from the native Ang II sequence.

Docked ACE2-peptide structures were obtained, where the peptide was coordinated to zinc in the ACE2 cleft as shown in Figure 2A. To be noted that the S1 binding region on ACE2, as obtained from literature^14^, is located outside the Ang II binding pocket. From the classification and known activity of ACE2, it is known that the zinc ion found in ACE2 participates in the cleavage of phenylalanine (F) residue from Ang II^10^. When Ang II peptide approaches the zinc ion located in the ACE2 cleft, ligand substitution can occur, with peptide replacing the water typically found coordinating to zinc. Several sites on Ang II can participate in this coordination including aspartic acid (D), tyrosine (Y), histidine (H), and the C-terminus of the peptide. Coordination with zinc at each of these sites can be quite strong, thus allowing a stable binding of the peptides to ACE2 without being cleaved, as was determined experimentally^13^. Four complexes were generated for each unique Ang II mutant, with a different residue of the peptide coordinated to zinc in each. Figure 2B-2E show the four structures generated for native Ang II, with Ang II coordinated to zinc at residues one (D), four (Y), six (H), and the C-terminus.

**Figure 2.**
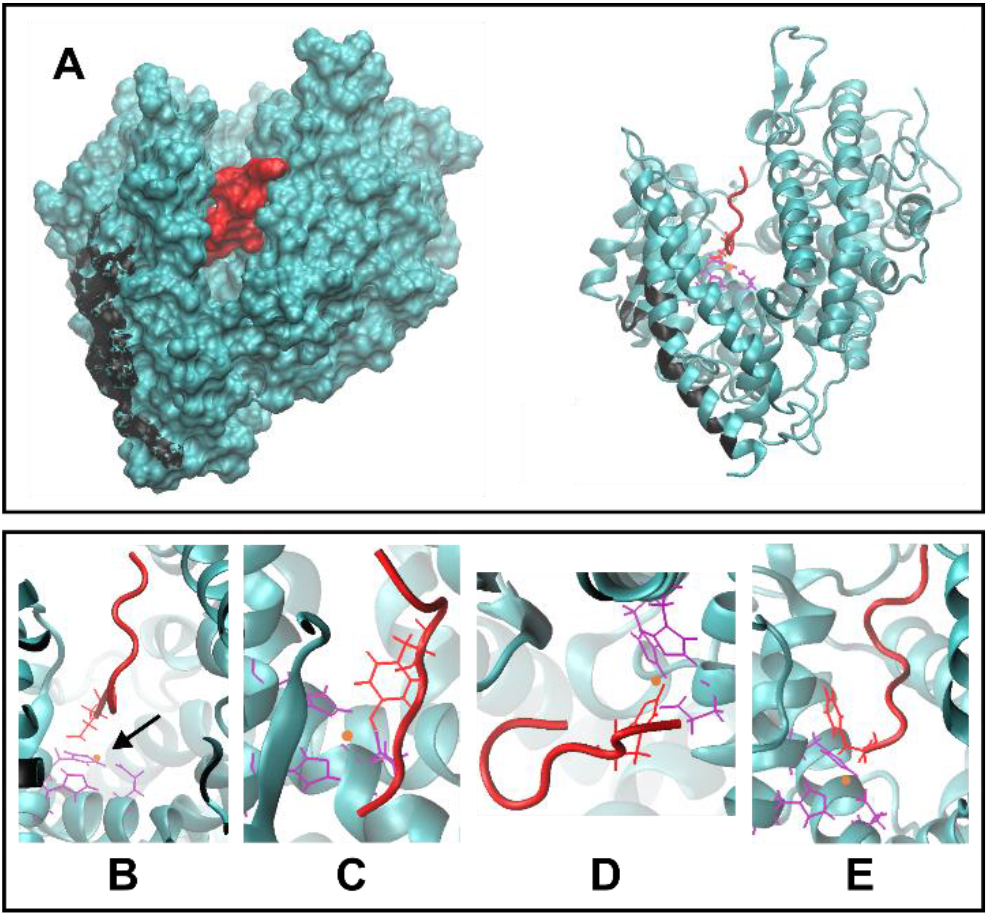
(A) The structure of Ang II (red) bound to ACE2 (teal). The cleft in ACE2 where Ang II resides is visible in both the surface-rendered image (left) and the ribbon diagram (right). The residues on ACE2 that interact with spike protein are shown in black. (B-E) Magnified views of Ang II-ACE2 binding, depicting the coordination of Ang II to zinc (orange) at four possible residues of D, Y, H, and the C-terminus, shown in B, C, D, and E, respectively. Corresponding coordinating residues on ACE2 are shown in purple.

Next, we examined the stability of the ACE2-peptide complex and compared the differences due to coordination at various sites of interest within the complex. The RMSD of the ACE2-peptide complexes was calculated throughout the duration of the 10 ns production run (Figure 3A). To verify that this runtime is sufficient to capture the true behavior, 50 ns production runs were performed for the native Ang II-ACE2 complexes (Figure 3A) to compare the effects of a longer simulation time. We found that the longer simulation time did not make a major impact on the average RMSD or magnitude of fluctuations, thus confirming the validation of the results from a shorter run of 10 ns. Interestingly, a noticeable difference was observed between the RMSD values of the whole ACE2-peptide complex (Figure 3A-B) and the peptide within the ACE2-peptide complex (Figure 3C-D). The RMSD values of a single peptide also demonstrated larger variability when amino acid residues at different positions of the chain were selected to coordinate with zinc.

**Figure 3.**
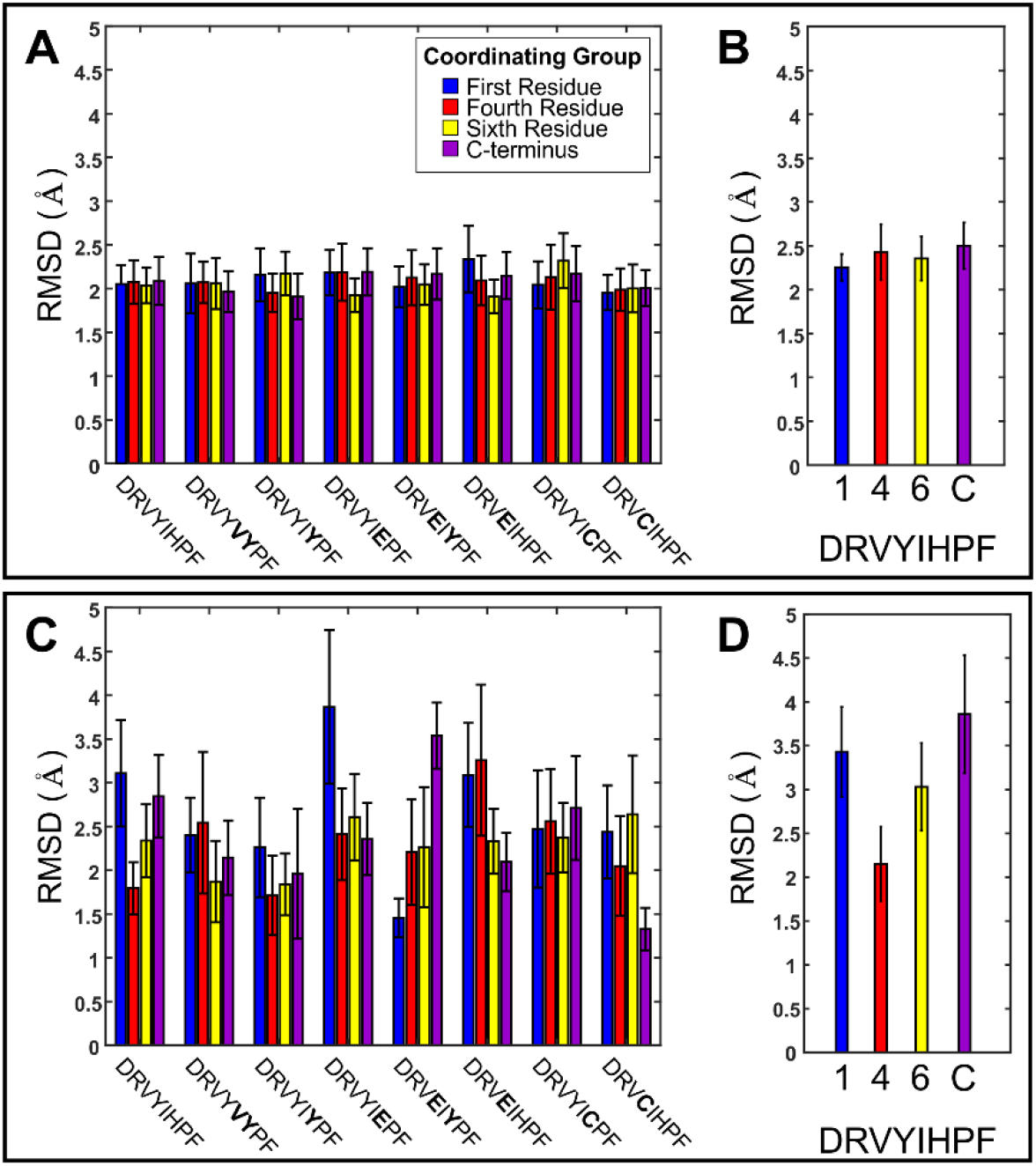
Computed averaged RMSD values for the entire Ang II-ACE2 protein complexes (A and B) and just the Ang II peptide or its mutant sequences (C and D). A and C correspond to 10 ns production runs while B and D correspond to 50 ns production runs. Each Ang II mutant has been coordinated to zinc at one of four possible locations on the peptide, as specified by bar color. The bolded residues in the peptide sequence are mutations from Ang II.

The variations in the binding site on ACE2 when native Ang II is coordinated to zinc at different amino acid residues are shown in Figure 4A-D. We found that beyond the movement of peptides during the production run, there were also some changes in the structure of the peptide binding site on ACE2. The changes in the secondary structure of the interacting residues, namely the residues within 3.5 Å of the peptide in the docked structure of ACE2, during the 10 ns production run is shown in Figure 4E-H. The binding free energy of each peptide, from the ACE2-peptide complexes, is plotted in Figure 5. While the interacting residues with α-helix and β-sheet secondary structures were relatively stable, fluctuations in structures were commonly observed for coils to turns. Furthermore, these changes were most prominent when the C-terminus of Ang II was coordinated to zinc.

**Figure 4.**
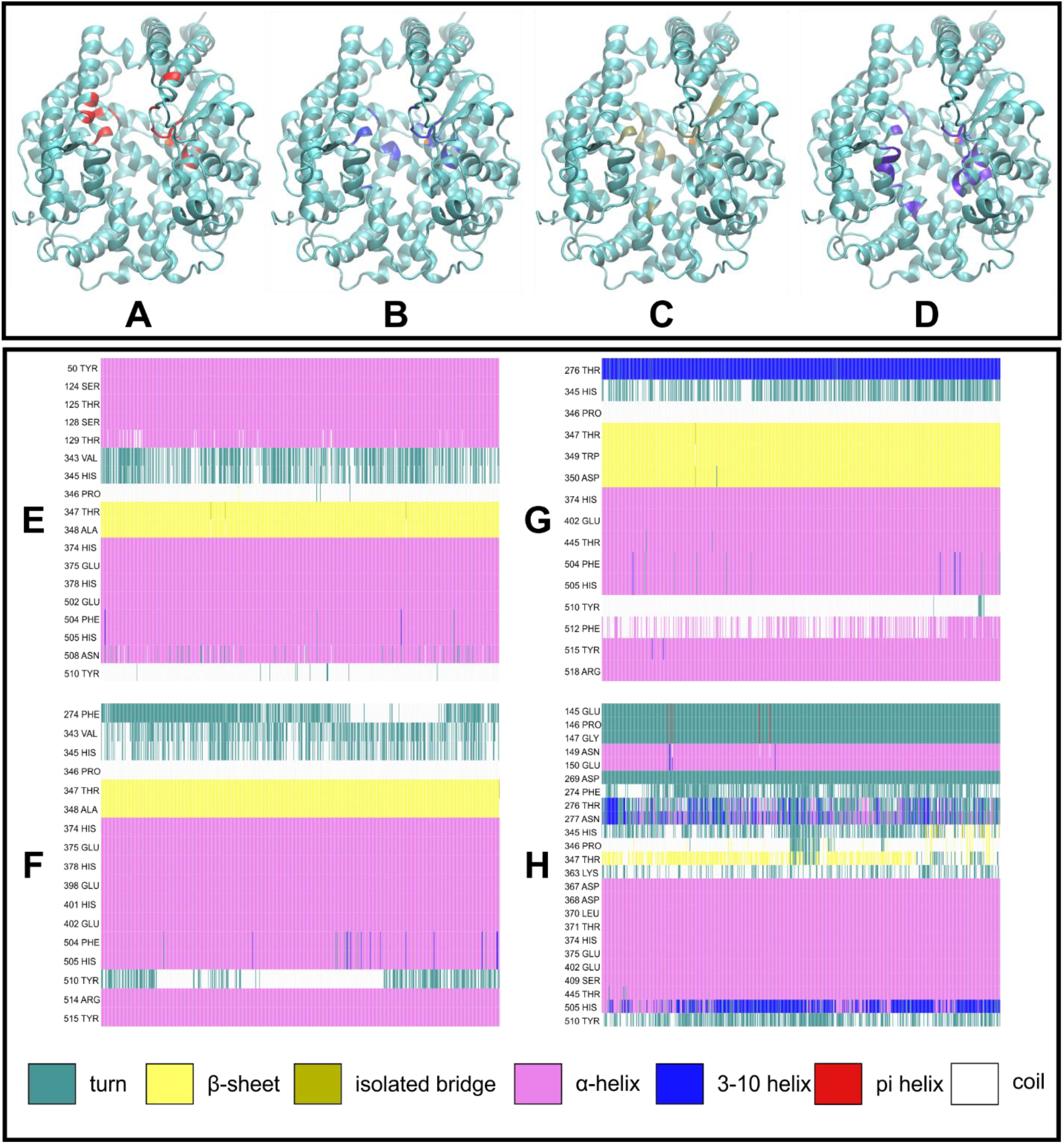
(A-D) ACE2 residues that interact with Ang II when coordinated to zinc at D, Y, H, and C-terminus respectively. Interacting residues of ACE2 are highlighted with a non-teal color. (E-H) Secondary structures of the interacting residues of ACE2 when Ang II is coordinated to zinc at D, Y, H, and C-terminus respectively throughout the 10 ns production run.

**Figure 5.**
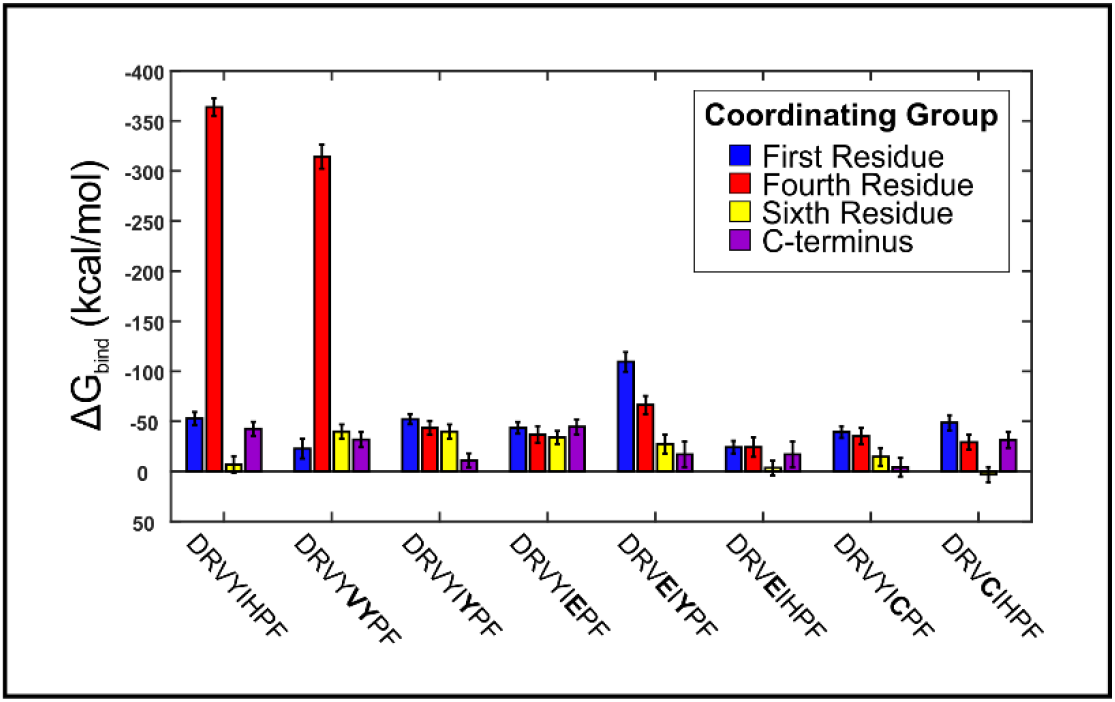
Binding free energy of native Ang II and mutant peptides found in the ACE2-peptide complexes. Residues at four different position in the peptide chain were considered for coordination to zinc. Bolded residues correspond to mutations from the native Ang II sequence.

The affinity of peptide binding to ACE2 was evaluated by calculation of binding free energy. The strongest binding free energies, with values more negative than -300 kcal/mol, were observed for native Ang II and DRVY**VY**PF mutant sequence when coordinated to zinc at the fourth residue (Figure 5). To further explore these unusually high values in comparison to other sequences, we made changes to the default MM/PBSA calculation. This modified approach took into consideration the energy contributions from changing zinc coordination bonds in the bonded model for zinc, as well as an adjusted free energy for unbound ACE2, which would have a different charge distribution on residues coordinated to zinc. After these adjustments, the resulting binding free energy of Ang II coordinated to zinc at tyrosine did not elicit any major change in the results, further suggesting there the observed difference is unlikely due to an artifact of the procedure for binding free energy calculation. The DRV**E**I**Y**PF sequence also showed a high binding energy, however, here the coordination to zinc was observed on the first residue (D).

Finally, we examined whether the interaction of Ang II or the mutant peptides could potentially elicit allosteric inhibition of S1 binding to ACE2 (Figure 6). The RMSDs of ACE2 residues at the S1 binding region, as identified by Han et al.^14^, were calculated during the production run of the ACE2-peptide complexes (Figure 6A). The average RMSD values in this figure were comparable to those of the entire protein during this simulation (Figure 3). Additionally, we studied the change in secondary structure of the interacting residues located at the S1 binding region during 10 ns production run (Figure 6B). While the α-helical structure remained stable during the run, some variability in the secondary structure at residues 82M and 355D were observed. However, this change is unlikely to impact the binding of spike protein to ACE2.

**Figure 6.**
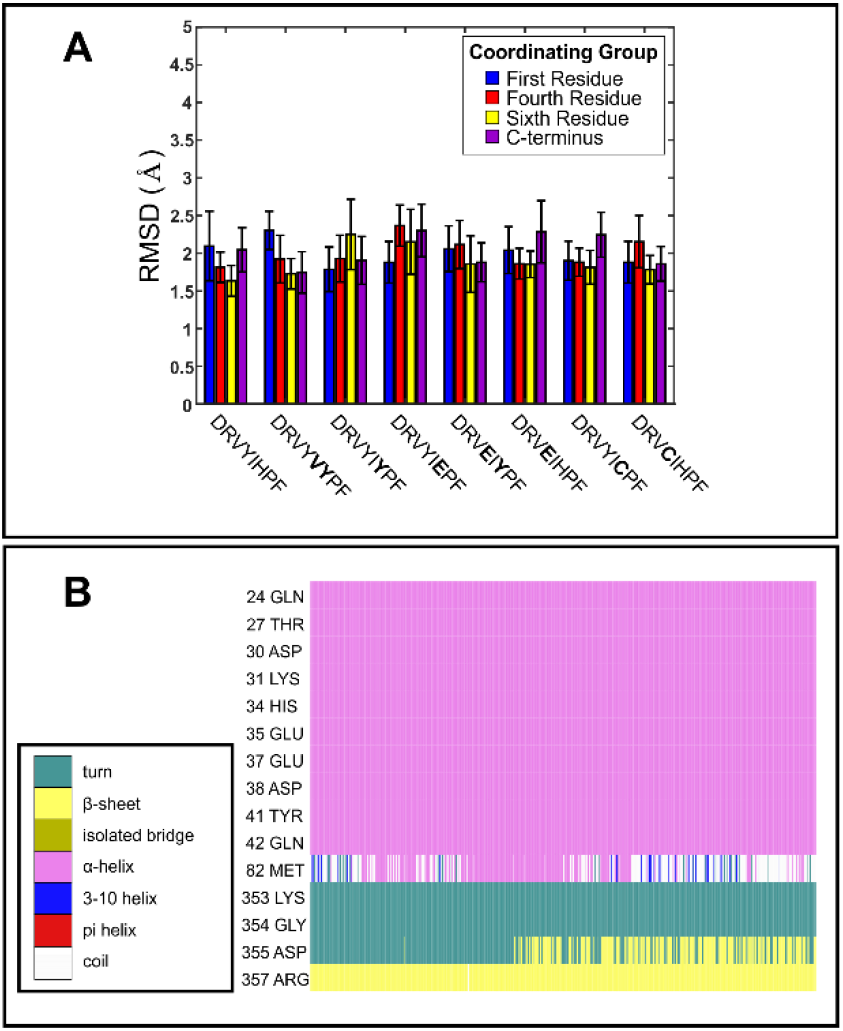
(A) Averaged RMSD of the S1 binding region on ACE2 during the production run of the various complexes with native Ang II and mutant sequences (mutations are shown in bold font). Each bar color represents a different residue location in the peptide chain coordinating to zinc in the complex. (B) Secondary structures of ACE2 residues that interact with spike protein throughout a 10 ns production run of Ang II-ACE2 complex with native Ang II coordinating to zinc at D. The box to the left explains how colors in the timeline plot correspond to secondary structure features.

## DISCUSSION

We found that binding of native and mutant Ang II peptides to ACE2 results in more rigid peptides along with a reorganization of the secondary structure of the binding residues in ACE2 cleft. While an increased peptide stability is expected to correspond with a strong binding energy, we observed strong binding energies for native Ang II and DRVY**VY**PF mutant. Lastly, it was found that Ang II binding to ACE2 might not be sufficient to affect the secondary structure of S1 binding residues of ACE2, suggesting that Ang II does not allosterically inhibit S protein-ACE2 interaction.

The production run for the ACE2-peptide complexes generated a higher average RMSD and standard deviation values for the peptides compared with the rest of the ACE2-peptide complexes. This difference can be attributed to the lack of strong secondary structure in the short peptides (Figure S3), allowing for greater flexibility and movement than the larger and more structured ACE2 protein. Our results also indicate that docking the peptides to ACE2 results in changes in the ACE2 secondary structure (Figure 4). While residues 274F, 343V, 345H, 363L and 510Y tend to alternate between the unstable turn and coil configurations in the native Ang II-ACE2 complexes, residues 129T and 512F become more stable as they shift from turns and coils to various helices and β-sheets.

The initial selection of mutants for this study was based on the experimental work reported by Clayton et al., which examined the peptides DRVY**VY**PF, and DRVYI**Y**PF.^13^ These two mutants were shown to exhibit a stronger inhibition of ACE2 than the native Ang II on a quenched fluorescent substrate assay and demonstrated a higher resistance to enzymatic cleavage. The results from this experimental work suggest that the 6^th^ residue of Ang II could play an important role in binding to ACE2. To further understand the role of amino acid residue located at this position, in this work, we investigated three additional mutant sequences DRVYI**E**PF, DRVYI**C**PF, and DRV**E**I**Y**PF. Specifically, we examined mutations with introduction of cysteine and glutamate as the 6^th^ residue since these amino acids are known to coordinate strongly to the zinc ion. Additionally, in our preliminary analysis on binding energy calculation of native Ang II, DRVY**VY**PF, and DRVYI**Y**PF, it was observed that coordination to Y at the fourth position resulted in strong binding to ACE2. To explore the potential role of 4^th^ residue in the binding event, we considered additional sequences with mutations at the fourth position (DRV**E**IHPF, DRV**C**IHPF, DRV**E**I**Y**PF).

Our analysis showed that native Ang II and DRVY**VY**PF each form a complex to ACE2 with a notably strong binding free energy more negative than -300 kcal/mol. The fourth amino acid residue (Y) of the sequences was found to coordinates with zinc in these situations. The reasons for the observed effects are not clearly understood at this time as tyrosine residue at fourth position did not result in similar strong binding free energies for other sequences considered. One potential limitation and a source of variability could be from the lack of entropy calculations in MM/PBSA protocol. However, these errors tend to be quite small, even with flexible ligands, and therefore, it is highly unlikely to make a major contribution to the differences found in our observations^55,56^. The potential role of kinetics also cannot be excluded as it plays a critical role in computing large binding free energies. It is possible that while these states are thermodynamically favorable, the kinetics presents a large activation energy barrier, preventing the formation of these final states. The impact of kinetics, which requires more computationally expensive techniques, however, is beyond the scope of the current work.

During the production run, the S1 binding region generated a similar RMSD to the rest of the protein complex, ruling out any increased movement in this region. Additionally, the secondary structure of this binding region remains largely unchanged except for the two residues at 82M and 355D. Both findings suggest very few structural changes in the S1 binding region in response to Ang II binding. Thus, a relatively little impact on the S1 binding region following ACE2-peptide interaction suggests that these peptides alone are less likely to allosterically inhibit the binding of SARS-CoV-2. Identifying a peptide sequence that provides a strong binding at the Ang II binding cleft on ACE2 could still be an important step toward developing novel therapeutic strategies to prevent SARS-CoV-2 infection. A high-affinity peptide sequence conjugated to a nanostructure will enable binding to the ACE2 proteins displayed on the cell membrane surface and can inhibit the interaction with the virus spike protein by physically blocking the access of the S1 binding domain. The efficacy of such strategy can further be enhanced by presenting the sequence at a very high density on a nanostructure such as self-assembled peptide nanofibers^32^, which would allow the additional advantage of multivalent binding^33^.

## CONCLUSIONS

We have performed molecular docking studies to examine the binding of native Ang II sequence and several mutant variants to ACE2 receptor, considering zinc coordination in the interaction. Among these sequences, we found that the native Ang II sequence is optimal for binding to the ACE2 cleft with only one of the mutant sequences demonstrating binding energies of similar amplitude. The strong binding energy was associated with a tyrosine residue in the fourth position of the peptide sequence coordinating to the zinc atom in the cleft. Furthermore, our analysis of S1 binding region suggests that native and mutant Ang II peptides alone are unlikely to inhibit the interaction between ACE2 and S protein. However, a knowledge of strong binding sequences for the ACE2 cleft could open the possibility of developing new strategies to prevent virus binding to cells. Specifically, presenting these sequences at a high density on a larger nanostructure could efficiently occupy the available ACE2 receptors on cell surface and physically block further access of the virus spike protein to the receptor binding site.

## METHODS

### Obtaining Structures

The experimentally derived structures for Ang II and ACE2, 1N9V^34^ and 1R42^9^ respectively, were obtained from the RCSB PDB (rcsb.org)^35^. Initial Ang II mutants’ structures were derived through use of PDB manipulator^36^ found in CHARMM-GUI^37–39^ to induce single-point mutations. To improve these structures, Visual Molecular Dynamics (VMD)^40^ was used to add a water box with a padding of 5 Å as well as add Na^+^ and Cl^-^ ions to neutralize charge and establish a NaCl concentration of 0.15 M. The mutants then underwent minimization under constant temperature (310 K) and pressure (1 atm) for 10 ps using Nanoscale Molecular Dynamics (NAMD)^41^. Before proceeding, Ramachandran plots created with MolProbity^42^ and RMSD plots created in VMD were analyzed to check the generated structures and the minimization procedure. The Ramachandran plots for each minimized peptide can be found in the SI Figure S4.

### Docking and Model Selection

To dock the desired ligands from Ang II or mutant peptides to zinc within the ACE2 structure, GM-DockZn^43^ was used. This program determines potential coordination sites on the ligand molecule and docks each site to the zinc ion. While models are typically sorted with a scoring function, in this scenario, models are preferably selected based on coordination geometry. It is critical that the coordination geometry is made accurate during docking, as it will become fixed during zinc parameterization using the bonded model. As previously demonstrated by Der, ideal coordination distances and angles involving zinc were used to score each docked model’s coordination geometry^44^. The values used for model evaluation are included in the SI Table S1. Using these scores, four structures were selected for each peptide sequence, coordinated to zinc at a different residue of the sequence.

### Zinc Parameterization

To prepare structures for parameterization, structural files were first converted from CHARMM to Amber format with Bio3D^45^ which includes the removal of all hydrogen atoms. The protonation state of each titratable residue was determined through version 3.2 of H++^46–48^, and hydrogens were added accordingly. To parameterize the zinc coordination site, MCPB.py^49^ was used. The bonded model was selected to reflect partial charges found on the various atoms within the coordination complex and prevent ligand substitution events during later MD simulations. During the MCPB.py procedure, bond and angle parameters for the zinc ion were generated using the Empirical method^50^. Restrained electrostatic potential (RESP) charges for atoms within the coordination complex were calculated using quantum mechanics (QM) simulations with GAMESS-US^51^. Using AmberTools20^52^, a water box with a padding of 10 Å and neutralizing ions were added.

### MD Simulations

Using the solvated and neutralized docked complexes, minimization was performed for 30 ps with a step size of 1 fs at 310 K and 1 atm. Equilibration of the docked complexes was then performed at 310 K with constant volume for 2 ns with a step size of 1 fs, followed by a 10 ns production run using the same conditions^53–55^. The calculations for the Ang II-ACE2 complexes were performed on Clarkson University’s ACRES cluster using the *Linux-x86_64-ibverbs-smp* version of NAMD. For the native and mutant Ang II sequences alone, production runs of 48 ns were performed, after 2 ns equilibration. The calculations of these relatively smaller sized systems were performed with a multicore version of NAMD on a Linux server.

### Binding Free Energy Calculation

The Molecular Mechanics/Poisson-Boltzmann Surface Area (MM/PBSA) approach was used to determine binding free energies utilizing trajectories created during the MD production runs. The CaFE^56^ plugin for VMD, which utilizes APBS^57^, NAMD, and VMD, was used to calculate the free energies of the ligand, receptor and docked complex. In this protocol, the free energy is computed for the receptor-ligand complex as well as the free ligand and receptor in each frame of the production run trajectory. NAMD is used to calculate the gas phase free energy while APBS and VMD calculate the polar and nonpolar solvation energies, respectively. The binding free energy is computed by subtracting the average free energies of the ligand and receptor from the average free energy of the receptor-ligand complex.

## ASSOCIATED CONTENT

### Supporting Information

The SI contains molecular structures, RMSDs, secondary structures, and Ramachandran Plots of mutant Ang II peptides as well as a table of ideal coordination geometries used in the screening process.

## ACKNOWLEDGEMENTS

Some of the computing for this project was performed on the ACRES cluster. We would like to thank Clarkson University and the Office of Information Technology for providing computational resources and support that contributed to these research results. Additional computational resources for this grant were provided by the National Science Foundation under Grant No. 1925596. The authors acknowledge use of the following simulation and visualization software packages: 1) NAMD and 2) VMD: NAMD and VMD, developed by the Theoretical and Computational Biophysics Group in the Beckman Institute for Advanced Science and Technology at the University of Illinois, Urbana-Champaign.

## Notes

### Competing Interest Statement

The authors have declared no competing interest.

### Summary of Updates

An error in the references has been corrected.

